# DaliScope: Distilling Biological Insights from Deep Structural Alignments

**DOI:** 10.64898/2026.11.09.765376

**Authors:** Hao Liu, Petri Törönen, Zhihan Zhang, Aleksi Laiho, Liisa Holm

## Abstract

Structural similarity searches reveal protein relationships that sequence alignments miss, but treating DALI results as isolated match lists misses deeper connections. We present DaliScope, an interactive Python framework that treats complete DALI target sets as structured populations for systematic analysis of protein structural neighbourhoods. DaliScope combines structural community detection with sequence profiling, Pfam annotation, residue mapping, and 3D visualization, enabling target sets to be subdivided and examined at the residue level.

We demonstrate two applications. In a multidomain protein with sparse domain annotation, structural analysis uncovered previously unannotated domains. A structurally anchored H×D … H signature occurred in 697 of 1,093 Pfam-annotated CUPIN domains and in 15 domains lacking Pfam annotation; relaxing the central residue constraint recovered 96% of Pfam-annotated CUPIN domains among the DALI targets. In a second application, a DALI search with transitive closure returned >12,000 hits. Applying an 80% coverage criterion to a conserved structural core reduced this set to 6,708 domain instances, resolved into 23 structural communities. (80% was chosen instead of the default 50% because the GOLD domain has a compact core with short loops.) Many communities were strongly enriched for individual Pfam families while providing structural context for 1,097 targets lacking Pfam assignments. Three-dimensional superimposition revealed structural continuity among related *β*-sandwich architectures through progressive changes in sheet geometry and auxiliary elements.

Together, these examples show how a single DALI query provides a structural anchor for organizing large neighbourhoods, extending interpretation beyond sequence-based annotations, and deriving structurally anchored sequence signatures from defined populations.

**Availability:** https://github.com/Liisa-Holm-group/DaliScope http://ekhidna2.biocenter.helsinki.fi/dali/colab.html

## Introduction

Structural similarity provides a powerful route to discovering relationships between proteins that may be difficult to recognize from sequence alone. In a typical structural query structure. This representation is effective for identifying individual related proteins, but it leaves much of the information contained in the complete search population unexplored. Once a search returns hundreds or thousands of structurally related proteins, the central question could change from *which proteins resemble my query?* to *what can the relationships among these proteins reveal about protein families, function, and structural organization?*

Systematic organization of protein space is well established, spanning curated structural taxonomies such as SCOP (Murzin et al., 1995) and CATH (Orengo et al., 1997), early automated fold classifications such as FSSP based on all-against-all DALI comparisons (Holm & Sander, 1994, 1996), and sequence profile-based family resources such as Pfam (Paysan-Lafosse et al., 2024) and SMART (Letunic & Bork, 2025). AlphaFold (Jumper et al., 2021; Varadi et al., 2023) and related approaches have now lifted much of the historical scarcity of structural data by providing models for a vast fraction of protein sequence space, allowing structural searches to reach far beyond the relatively small set of experimentally characterized structures. This expansion has shifted the emphasis toward structure-aware approaches that can efficiently search and organize increasingly large structural datasets, including compact structural representations such as structural alphabets and protein language model embeddings (van Kempen et al., 2024; Heinzinger et al., 2024; Liu et al., 2025). Recent methods have also demonstrated multiple-structure alignment at very large scale, extending structural correspondence to collections containing hundreds of thousands of protein structures (Gilchrist et al., 2026). DaliScope addresses a complementary problem: how to organize and interpret the structural relationships within the complete target population returned by a query-centred DALI search.

DALI derives query-anchored multiple-structure alignments (MSTAs) directly from Cα coordinates and provides query-anchored transformations that place related structures in a common coordinate frame. DaliScope exploits this property to recover relationships within a large DALI neighbourhood without exhaustive all-against-all comparison, showing that a single query can provide sufficient structural context to organize the relationships among its many targets. Although DaliScope is not intrinsically tied to how an MSTA is generated, DALI provides a particularly suitable input through its high-quality structural alignments (Holm, 2020; Chen et al., 2025).

DaliScope treats the complete DALI target set as a structured population rather than a ranked list of independent hits. The MSTA provides the fixed structural correspondence from which population-specific structural profiles, sequence profiles, and residue-level signatures can be derived, while spatial fingerprinting and pairwise geometric scoring organize the target set into structural communities. These populations can then be examined through three-dimensional structural mapping and visualization. Thus, rather than requiring a global classification of protein space, DaliScope uses a single focal query to model the structural neighbourhood around it, organize that neighbourhood into populations, and derive sequence and structural signatures from the resulting structural context.

## Methods

### DaliScope data model and workflow

DaliScope operates on the complete target set returned by a DALI structural similarity search (Holm, 2022; Liu et al., 2025). A self-contained data pack contains the DALI alignment statistics, the DALI multiple-structure alignment (MSTA), Pfam 38.2 annotations, and Cα coordinates required for downstream analysis. DaliScope provides interactive operations for selecting targets, restricting alignment positions, generating analytical subsets, and visualizing their sequence and structural properties. Detailed implementation and parameter descriptions are provided in the Supplementary Methods.

We use two concepts throughout the DaliScope workflow. A population is a defined subset of the DALI target set that is carried forward for analysis. A population may comprise the complete target set, a user-selected subset, or a subset generated by an analytical procedure, such as a structural community or sequence-defined subfamily. A view is a representation of a population in which the targets and MSTA positions relevant to a particular analysis have been selected. Thus, populations define which targets are being analyzed, whereas views define which parts of the available structural and alignment data are being examined. For example, domain clipping generates a derived view in which the superimposition is updated.

### DALI reference alignment and fold-space representation

DALI provides the initial multiple structural alignment (MSTA) correspondences and anchor-derived coordinate transformations used to place all targets in a common reference frame. These residue equivalences serve as the basis for profile logos, residue mapping, and visualization in DaliScope. However, to isolate rigid structural modules, domain clipping applies occupancy weighting that assigns zero weight to non-coherent or peripheral MSTA column correspondences. This selective zero-weighting suppresses spatial noise from mobile or non-rigid segments without altering the underlying MSTA coordinate frame.

To construct similarity networks and evaluate structural continuity within the target population, the Rigid Core extension requires pairwise correspondences that can extend beyond the bounds of the clipped MSTA. For each candidate target pair, residue equivalences are recomputed *de novo* from coordinates expressed in the DALI-derived reference frame using a dynamic-programming procedure based on intermolecular (C_\alpha) distances. This yields an optimized, sequential, non-crossing pairwise alignment used exclusively to calculate inter-target edge weights, while leaving the global MSTA topology intact.

Structural similarity and alignment coverage are mapped alongside Pfam domain and clan annotations to represent the structural organization of the target set.

Interactive views enable filtering and inspection of target populations at progressively finer resolution, providing a systematic starting point for identifying structural domains, multidomain architectures, and conserved functional signatures.

### Structural community detection

Structural similarity can extend beyond the region represented by the query, allowing target structures to share additional structural elements that are absent from the query itself. To identify such relationships and organize them into structural communities, we developed a voxel-based fingerprinting and pairwise geometric rescoring pipeline followed by network community detection. DALI-derived superimpositions provide the common structural reference frame, but the similarities used for structural clustering are computed independently of the DALI alignment.

The pipeline first uses sparse, IDF-weighted voxel fingerprints of Cα traces to rapidly retrieve approximate structural neighbours. Retrieved pairs are then rescored using a geometric similarity measure based on intermolecular Cα distances, with residue correspondences determined *de novo* and resolved into a sequential, non-crossing alignment by dynamic programming.

The voxel-based candidate pre-filter restricted pairwise geometric scoring to 200 candidate neighbours per target. This candidate-set size was calibrated against exhaustive pairwise scoring; the 200 retrieved candidates contained >95% of the top-30 neighbours identified by exhaustive scoring (Supplementary Methods). The resulting distance-weighted similarity scores are assembled into a sparse nearest-neighbour network and partitioned into structural communities using Infomap (Rosvall & Bergstrom, 2008). Module distinctiveness and cohesiveness are evaluated from within- and between-module similarity distributions and by recursive subdivision, while correspondence with Pfam classification provides an independent assessment of module organization.

Each resulting structural community is treated as a new population within DaliScope. It can therefore be filtered, visualized, aligned, and analyzed using the same downstream workflows as the original target set.

### Sequence profiling and structural interpretation

DaliScope generates sequence profiles for selected populations using the fixed DALI MSTA. Because the sequence positions retain their correspondence to the DALI structural alignment, profile positions can be mapped directly to residue positions in the corresponding structures. Interactive sequence and structure views synchronize alignment positions with three-dimensional structural representations. This enables conserved or discriminatory positions identified in a sequence profile to be examined directly in their structural context.

### Signature Motif Pipeline

The Signature Motif Pipeline derives and tests sparse sequence signatures associated with selected structural or functional subfamilies. A seed population is examined through its DALI MSTA to identify informative alignment positions, which are selected to define a Prosite-like (Sigrist et al. 2002), position-specific signature anchored to the corresponding MSTA positions. The resulting signature is applied to the broader target population to identify and score sequences matching the specified residues at these structurally equivalent positions.

Signature-positive targets are returned as a new population that can be inspected using the same sequence, structural, and annotation views. Iterative comparison of these populations allows signatures to be refined and used to define sequence subfamilies that may not be separable by global structural similarity alone.

### Integrated population analysis

DaliScope links structural, sequence, and annotation-based analyses through the population/view framework. Structural communities generated from the similarity network become populations for sequence profiling and motif analysis; conversely, sequence-defined populations can be projected onto the structural network and examined in three-dimensional context. This iterative subdivision of the original DALI target set enables analysis to proceed from global fold similarity through structural communities to increasingly specific sequence-defined subfamilies.

## Availability

The DaliScope frontend generates a self-contained data pack from a DALI server result set and launches the DaliScope environment in Google Colab: http://ekhidn2.biocenter.helsinki.fi/dali/colab.html Source code is available on GitHub: https://github.com/Liisa-Holm-group/DaliScope

## Results

### Filling gaps in domain family annotations

Full details of this analysis are provided in Worked Example 1 (Supplementary Text ST1). We selected the AlphaFold (Jumper et al., 2021) model of rat lysine demethylase 3B (RAT:AF-M0RDF1-F1) as a challenging query because its 1,722-residue sequence has sparse Pfam annotation (Figure 1). A Dali web server search of the AlphaFold Database v.2 (Varadi et al., 2023) returned 1,573 hits. By visual inspection, the query structure contains five compact domains at residues 1–149, 150–214, 840–920, 921–1100, and 1343–1722 that correspond to five distinct, non-overlapping blocks in the DALI MSTA. The intervening segments are modelled with low confidence by AlphaFold.

**Figure 1:**
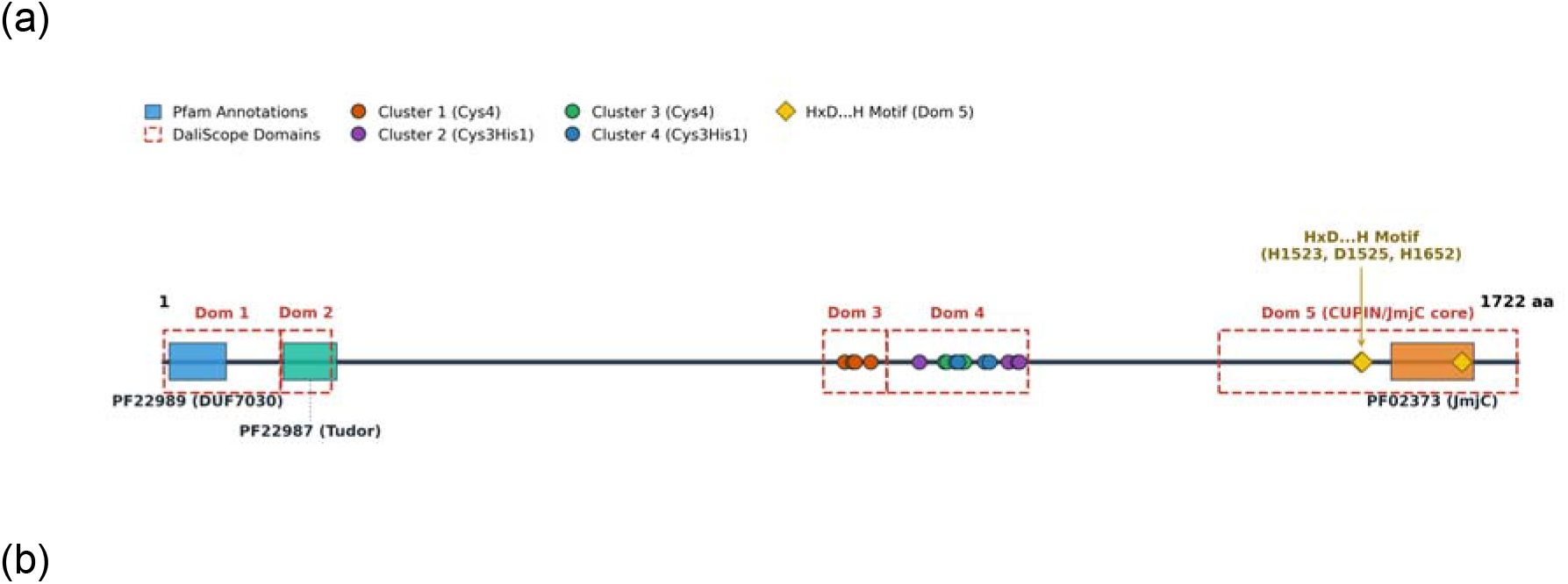
Worked Example 1 – comparing structural to sequence based annotation. Domain architecture of rat KDM3B (query sequence, 1,722 aa). Known Pfam domains are indicated by solid filled boxes, while structural domains refined in this study are shown as open dashed red boxes. Candidate metal-binding clusters (Domains 3 and 4) are marked by coloured dots, and the catalytic HxD…H motif (Domain 5) is indicated by filled diamonds (HxD diamonds overlap).

Population analysis showed that these domains differ markedly in their structural and sequence associations. Domains 1 and 2 adopt SH3-like folds, and occur in many proteins that lack Domains 3–5 (9/16 and ∼108/111 targets, respectively).Domains 3–5 formed distinct structural populations within the broader target set, with strong associations to CUPIN domains. Several instances of Domains 1 and 2 lacked Pfam annotation. Domains 3 and 4 showed distinct associations with CUPIN domains that were not captured by current Pfam models, while partial occupancy of Domain 5 indicated that its structurally conserved core extends beyond the corresponding Pfam sequence profile.

Sequence logos (Tareen & Kinney, 2020) of the structurally defined populations revealed conserved cysteines and histidines in Domains 3 and 4. Mapping these conserved positions onto the query structure identified four spatial clusters with Cys4 or Cys3His1 arrangements, consistent with candidate metal-binding sites. Domain 3 had no Pfam-annotated targets in the DALI data. Four of 51 targets containing Domain 4 had Pfam annotations to a zinc finger domain of monoamine-oxidase A repressor R1 (PF10497, Chen et al., 2005). Pfam reports PF10497 in 156 distinct domain architectures, including several that involve Jumanji/JmjC domains. Domain 5 is a demethylase annotated as a member of the 2OG-Fe(II) oxygenase superfamily (PF03171) of the CUPIN clan (CL0029).

Sequence logos recovered the known 2OG-oxygenase/JmjC motif HxD…H in Domain 5 (Balciunas & Ronne, 2000). In DaliScope, this sparse sequence motif is anchored to the DALI MSTA, allowing it to be evaluated at structurally corresponding residue positions across the full target population of 1,176 structures, of which 1,093 were annotated as CUPINs. Unlike an unconstrained sequence search, this approach therefore evaluates the motif within the structural correspondence defined by the MSTA. The strict HxD…H configuration occurred in 697 of the 1,093 annotated CUPINs, predominantly among dioxygenases, and was also present in 15 previously unassigned domains. The broader CUPIN superfamily is functionally diverse, so variation in the configuration of residues contributing to the active site is expected.

Consistent with this, residue-frequency analysis showed that the central position of the motif was variable. Relaxing the signature to require only the two conserved flanking histidines recovered 96% of the annotated CUPINs in the dataset, extending the structurally anchored signature beyond the dioxygenase-dominated subset to encompass most of the broader CUPIN population.

Together, these analyses show how DaliScope transforms a large DALI neighbourhood from a query-centred set of structural matches into a population-level representation of fold organization. DaliScope delineated five recurrent structural domains, identified previously uncharacterized CUPIN-associated structural regions, revealed conserved sequence and structural features including candidate metal-binding sites, and enabled a structurally anchored motif search across the population. Using Pfam clan assignments as an independent reference, we showed that sparse, structurally anchored sequence motifs could recover nearly the entire CUPIN superfamily from a single DALI target set, providing a complementary route to superfamily-level organization that does not require the multiple family-specific sequence profiles used by Pfam. The resulting analysis provides a framework for interpreting sequence features that are difficult to identify from primary-sequence annotation alone.

### Fold-space continuity

Our second worked example (Supplementary Text ST2) addresses a complementary problem: how to interpret a large structural neighbourhood when a single query is similar to multiple domain families and superfamilies. GOLD-like domains (Pfam clan CL0521) resemble CUPIN domains; both are small β-sandwich structures with a jelly-roll topology. We chose a single-domain query from human Golgi resident protein GCP60 (PDB 5TDQ, chain A). The initial DALI search with transitive closure with default parameters (Liu et al., 2025) of the AlphaFold Database v.2 (Varadi et al., 2023) returned more than 12,000 structurally related hits.

From this target set, we defined a coherent structural core from secondary-structure elements. The resulting MSTA core spanned residues 23–123. Applying an 80% coverage criterion reduced the initial target set to approximately 6,708 domain instances while excluding fragmentary matches. The resulting population was defined directly from structural correspondence rather than external domain annotations.

The structural network resolved the domain-fold population into 23 communities, many of which showed strong enrichment for individual Pfam families (Figure 2). Because Pfam annotations were not used to construct the network, this correspondence provides an independent comparison between structural and sequence-based classification. The communities also included many targets without Pfam assignments, including approximately 1,000 in the unassigned category. These targets were not structurally isolated but occupied coherent regions of the network alongside annotated domains, providing structural context where sequence-based classification was absent.

**Figure 2:**
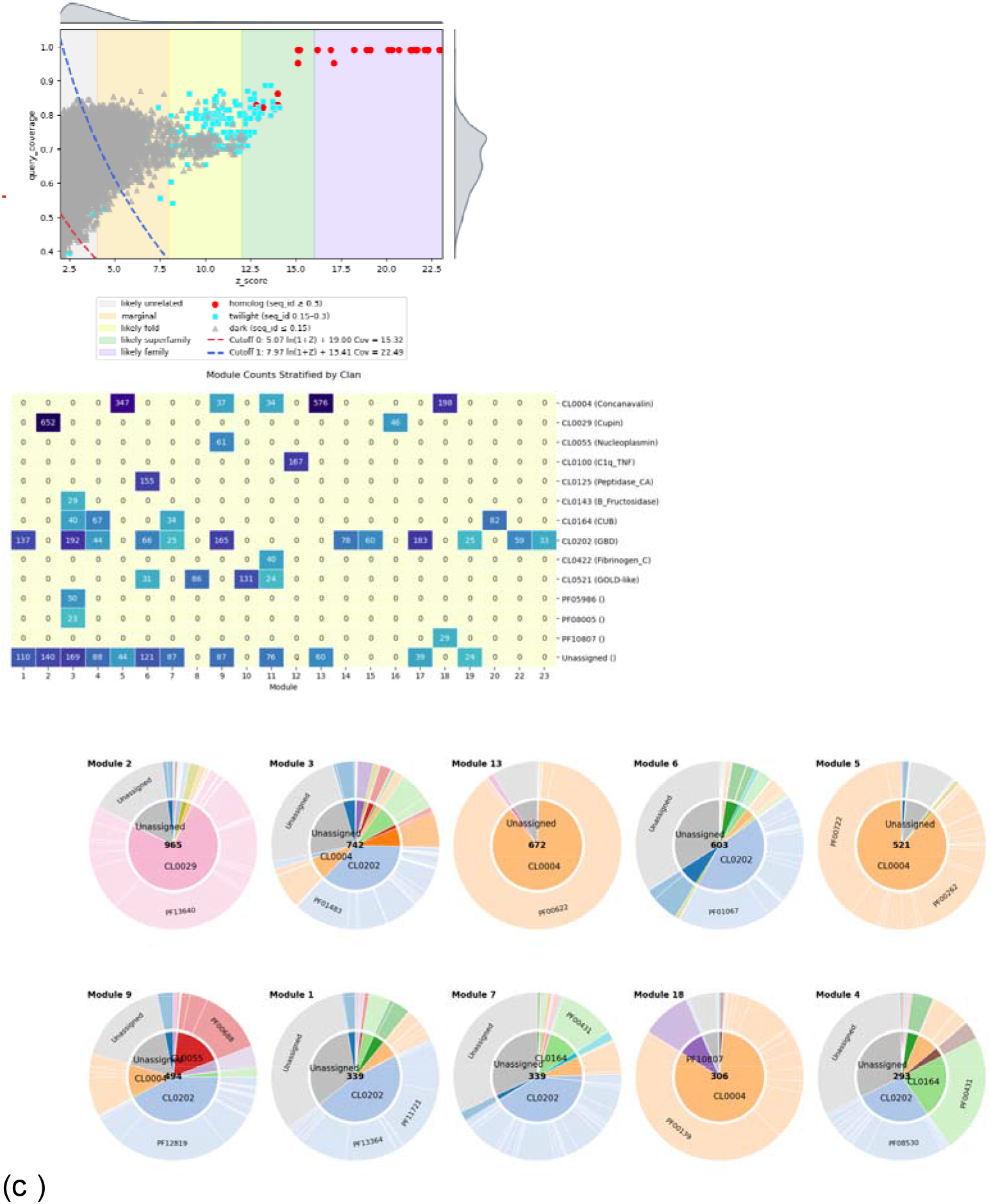
Worked Example 2 – Rigid Core Pipeline resolves relationships between DALI hits. **(a)** Overall distribution of DALI target hits in the (Z-score, query coverage) plane. The query protein has close homologs among the targets (red), while the full data set includes a mixture of remotely related Pfam superfamilies. Heuristic Z-score bands and cuts at valleys of the density distribution are shown for orientation, using default heuristic thresholds. **(b)** Structural community detection resolves fold-space continuity into coherent structural modules enriched with particular clans. Counts <10 are not shown. Note that Pfam annotations were not used during community detection. **(c)** Sunburst diagrams illustrate the Pfam family composition of the ten largest communities (module sizes ranging from 293 to 965 targets), coloured by Pfam clan. Gray sectors denote targets unassigned by Pfam.

Recursive community detection revealed that fold-space structure occurs at multiple scales. Most communities remained cohesive under recursive subdivision, whereas some separated into distinct subcommunities, for example a subdivision distinguishing PF17065 from the Concanavalin-like CL0004 group. Consistent with this organization, within-community pairwise similarity scores were higher than between-community scores, supporting the structural coherence of the detected communities.

The DALI-derived transformations placed all targets in a common coordinate frame anchored on the conserved domain, allowing the shared core to be superimposed while peripheral structural variation remained visible. Thus, the query provided a structural anchor for visualizing continuity across a much larger fold space. Three-dimensional superimposition revealed that the boundaries between communities did not necessarily correspond to abrupt changes in the underlying fold. Related β-sandwich architectures could instead be connected through progressive changes in sheet geometry, including the addition or loss of edge β-strands, changes in strand connectivity, and the emergence of auxiliary structural elements (Figure 3).

**Figure 3:**
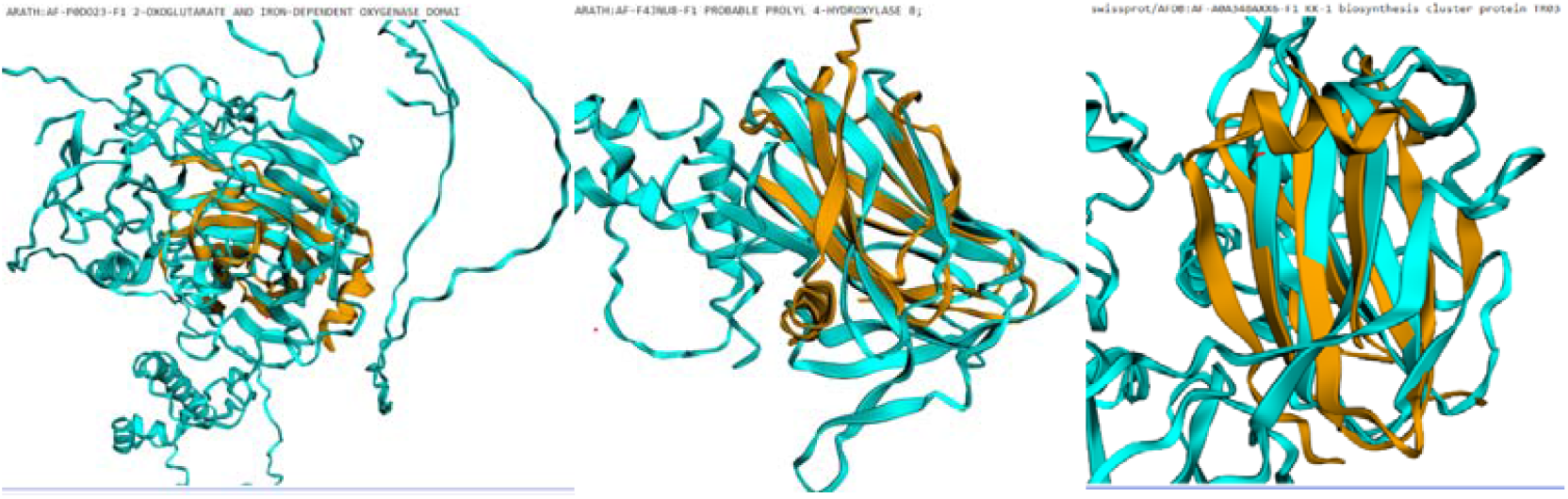
Structural variations. Examples of similar folds superimposed onto the GOLD-like domain used as query representing, from left to right, CUPIN, PF25238, and a Pfam-unassigned target. The jellyroll topology of the core is shared, but strands are elongated, bulged, and added or deleted at the edges. Additional mass can pack against the beta-sandwich. The Rigid Core Pipeline was designed to capture the conservation of such auxiliary structural features.

Together, these analyses show how DaliScope extends a large DALI neighbourhood from a query-centred set of structural matches into a population-level representation of fold organization. Structural communities can be compared with independent sequence-based classifications, while Pfam-unassigned targets can be placed within coherent structural populations. Recursive subdivision and three-dimensional inspection further characterize how structural similarity is organized within and between these populations. Whereas Worked Example 1 uses structural alignment to resolve residue-level features and hidden domains within a single multidomain protein, this example applies the same structural information at population scale to characterize the organization, subdivision, and continuity of an extended fold space.

### Computational performance

For the Worked Example 2 dataset, voxel-based candidate indexing required 1.00 s, while subsequent geometric scoring of 1,341,600 candidate pairs required 73.34 s on a single CPU thread, corresponding to an average of 54.67 μs per pairwise evaluation. Because the candidate-neighbourhood size was fixed, the number of pairwise geometric evaluations in the rescoring stage scaled approximately linearly with the number of targets, rather than quadratically as in exhaustive all-against-all comparison.

## Discussion

DaliScope changes the primary unit of analysis in structural similarity searching from the individual hit to the target population. A conventional DALI search primarily answers which structures are similar to a query and ranks those relationships by pairwise score. The results presented here demonstrate that the target set itself contains rich, actionable biological information: targets can be organized into structural communities, cross-referenced with sequence annotations, and mined for recurring residue-level signatures. Treating the DALI neighbourhood as a population provides an intermediate level of analysis between pairwise structural comparison and global classification, enabling relationships among hundreds or thousands of structurally related targets to be interpreted within a unified framework Worked Example 1 illustrates the value of this population view for incomplete domain annotation. Structural populations derived from a single multidomain query exposed recurrent regions that were only sparsely represented by Pfam annotations and provided a structural context in which conserved residues could be identified and mapped. The CUPIN analysis further showed that a sparse sequence motif could be evaluated at structurally corresponding positions across more than 1,000 annotated CUPIN targets, distinguishing a more restrictive dioxygenase-associated configuration from a broader population-level signature. Thus, structural correspondence can provide a coordinate system for sequence interpretation even when sequence-profile annotation is incomplete or when a family is too diverse for a single short sequence pattern to capture its full population.

Worked Example 2 demonstrates a complementary consequence of the population-centred approach. The query did not need to encompass the full structural diversity of the resulting neighbourhood for that neighbourhood to be informative. DALI-derived transformations provided a common structural reference frame, while independently recomputed pairwise structural similarities organized the target population into communities. These communities showed correspondence with Pfam families despite Pfam annotations not being used to construct the network, and targets lacking Pfam assignments could be interpreted in the context of neighbouring annotated populations. Three-dimensional inspection further showed that community boundaries could occur within a continuum of related β-sandwich architectures rather than at discrete structural discontinuities. This illustrates how a query-centred structural neighbourhood can provide a local representation of fold organization without requiring a global classification of protein space.

DaliScope is designed to complement, rather than replace, sequence-based annotations and global classification systems. While Pfam provides family-specific sequence profiles and curated assignments, DaliScope uses structural correspondence to organize the target population returned by a search and uncover localized sequence features within that structural context. Similarly, DaliScope does not seek to replace global databases (such as SCOP or CATH) or global MSTA algorithms; its goal is specifically to exploit the internal organization of a query-centered neighbourhood after a search is executed. This distinction is critical: the populations examined by DaliScope are bounded by the structural perspective of a particular query and need not strictly conform to globally delimited evolutionary families.

Several limitations follow from this architecture. DaliScope remains dependent on the quality and representation of the underlying DALI search and MSTA, and its structural communities should not be conflated with formal phylogenetic or evolutionary trees. Additionally, network edge construction relies on voxel-based candidate retrieval followed by pairwise geometric rescoring rather than exhaustive all-against-all alignment; while candidate-set calibration provides an empirical measure of retrieval recall, the network is an approximation of an exhaustive graph. Likewise, the fidelity of structurally anchored sequence signatures depends directly on alignment coverage across the population.

As structural databases continue to expand through both experimental determination and accurate structure prediction, the number of meaningful relationships returned by a single structural search will continue to increase. DaliScope provides a way to retain that population-level information rather than reducing the search to a ranked list of pairwise similarities. By combining structural organization with sequence profiling, residue mapping, annotation and three-dimensional inspection, it provides a route from a query-centred structural neighbourhood to testable structural and sequence hypotheses.

## Supplementary Materials

Supplementary Methods (Word doc)

ST1: WorkedExample1.pdf (Notebook outputs)

ST2: WorkedExample2.pdf (Notebook outputs)

### Declaration of generative AI and AI-assisted technologies

AI-based tools were used to assist with language editing and polishing of the manuscript and with development and refinement of computer code. All scientific content, analyses, interpretations, and final text were reviewed and verified by the authors, who take full responsibility for the content of the manuscript.

