## Supplementary Methods for "DaliScope: Distilling Biological Insights from Deep Structural Alignments"

**S1. DaliScope data model, populations, and views**

DaliScope analyses the complete target set returned by a DALI structural similarity search. The input is distributed as a self-contained data pack containing DALI alignment statistics, the DALI multiple-structure alignment (MSTA), Pfam annotations, and Cα coordinates required for downstream analysis.

DaliScope uses two concepts throughout the workflow: populations and views. A population is a defined set of targets derived from the original DALI target set. The initial DALI target set constitutes the starting population; filtering, structural community detection, or sequence-signature analysis can generate additional populations.

A view is a representation of a population in which targets and MSTA positions have been selected for a particular analysis. Row selection determines the targets represented by the view, whereas column selection restricts the MSTA to the structural or sequence region under investigation. Domain clipping generates a derived view in which the structural superimposition is updated.

**S2. DALI MSTA and structural coordinate framework**

DALI provides the initial structural alignment, the residue correspondence used throughout sequence-based analyses, and rigid-body transformations for placing target structures in a common query-anchored coordinate frame. For each target, the transformation obtained from the DALI domain-anchored alignment was applied to its Cα coordinates,

$$x^{'}=xR^{T}+t,$$

where $R$and $t$are the rotation and translation obtained from the DALI alignment. The transformation is determined using the user-specified query/anchor domain and then applied to the full target structure, thereby placing all targets in a common coordinate frame.

The **DALI MSTA** is retained as the fixed residue correspondence throughout DaliScope. Sequence profiles, MSTA displays, structural-region selection, and mapping between alignment positions and structure residues all use this correspondence. No new sequence correspondence is introduced during these analyses.

The DALI-derived coordinate frame is used for coordinate-based visualization and voxel fingerprinting. It provides a shared spatial reference for comparing target geometries, but does not define the pairwise structural similarity used for community detection. Candidate target pairs are instead evaluated independently using a geometric similarity measure based on intermolecular Cα distances, with residue correspondences determined de novo for each pair. Thus, the MSTA defines the fixed residue correspondence, the DALI transformations define the common coordinate frame, and the pairwise geometric score provides the target–target similarity used to construct the structural network.

**S3. Domain clipping and refined superimposition**

DaliScope can refine a structural population by selecting a region of the DALI MSTA and restricting the analysis to targets that sufficiently cover that region. Structural regions are selected by restricting a view to specified MSTA columns. A coverage criterion can then be applied to retain only targets covering a specified fraction of the selected region.

For a selected population and structural region, DaliScope performs an occupancy-weighted structural refinement. For each MSTA position, the occupancy weight is proportional to the frequency with which that position contains an aligned residue among members of the population. All-gap columns are excluded. Positions consistently represented across the population therefore contribute more strongly to the refined superimposition, whereas sparsely represented positions contribute less.

The resulting population-specific superimposition provides a structural representation of the region shared by the selected population. It is used for population characterization, structural visualization, and the Rigid Core Pipeline. This refinement does not alter the DALI MSTA correspondence or the pairwise similarity scores used to construct the structural community network.

When domain clipping is applied, the selected MSTA region and its coverage-filtered targets define a derived population-specific structural view, for which the superimposition is recalculated. This allows recurrent structural regions to be analyzed independently of existing sequence-family boundaries, including regions that extend beyond or are not represented by existing Pfam models.

**S4. Voxel-based structural fingerprinting**

Voxel fingerprinting provides a computationally efficient pre-filter for identifying candidate structural neighbours. It is a coarse candidate-generation procedure and is not used as the final measure of pairwise structural similarity.

For each target in a population, Cα coordinates in the common reference frame were discretized onto a regular cubic grid with a voxel edge length of 4.5 Å. Each occupied voxel was recorded once per target as a binary target-level occupancy, together with a dwell count corresponding to the number of residues mapping to that voxel. Where specified, residues belonging to the query/anchor domain can be excluded from the fingerprint to emphasize accessory structure.

An inverse-document-frequency weight was assigned to each voxel,

$$\mathrm{IDF}\left( v \right)=\min\left[ \log\left( \frac{N+\alpha}{n_{v}+\alpha} \right) , c \right],$$

where $N$is the number of targets in the population, $n_{v}$is the number of distinct targets occupying voxel $v$, $\alpha$is the additive smoothing constant, and $c$is the upper weight cap. The resulting fingerprint combines voxel dwell count and IDF weighting. Where a query-domain reference point is used, a radial damping term,

$$D\left( r \right)=\frac{1}{1+\left( r/r_{0} \right)^{2}},$$

with $r_{0}=20$Å, can additionally reduce the contribution of voxels distant from the query-domain centre of mass.

To reduce sensitivity to small coordinate displacements at voxel boundaries, occupied voxels are expanded using a local matching kernel. The default kernel is a seven-point kernel consisting of the occupied voxel and its six face-adjacent neighbours, with adjacent voxels assigned a relative weight of 0.25.

An inverted index stores voxel identities together with target identities and fingerprint weights. For each query target, kernel-expanded voxels are used to retrieve candidate targets from the index. Soft voxel-overlap scores are accumulated for each candidate and normalized by the L2 norm of the corresponding raw, unexpanded fingerprint. The top $k_{2}=200$candidates for each target are retained for independent pairwise geometric rescoring.

The voxel score is therefore used only for candidate retrieval and does not define the the pairwise structural similarity used to construct the structural network.

**S5. Pairwise geometric alignment and rescoring**

Candidate pairs identified by voxel fingerprinting are rescored independently of DALI's MSTA residue equivalences while retaining the DALI-derived reference coordinate frame. The pairwise geometric similarity score is calculated *de novo* as an absolute, unnormalized sum over spatially matching $C_{\alpha}$ positions. Unnormalized scoring prioritizes total structural overlap, ensuring that significant local similarities in multidomain proteins are not suppressed by large, unaligned peripheral regions.

For each candidate pair, spatially compatible Cα residue pairs are identified using a k-d tree search with a distance cutoff of (2$d_{0}$), where ($d_{0}$=5.0) Å. Each candidate correspondence ( (a,b) ) receives the distance-dependent score

$$s\left( a , b \right)=\frac{1}{1+\left( d_{ab}/d_{0} \right)^{2}},$$

where $d_{ab}$ is the Cα–Cα distance between the two residues.

Candidate correspondences are resolved into a single sequential, non-crossing alignment. Residue indices must increase simultaneously along both structures, excluding crossing or unordered correspondences. The optimal correspondence chain is obtained by dynamic programming over the candidate pairs using a Fenwick-tree-accelerated prefix-maximum calculation (Fenwick, 1994). This permits the aligned region to extend beyond the original DALI anchor domain wherever supported by the pairwise geometry.

The pairwise geometric similarity score is the sum of the distance-weighted correspondence scores along the optimal chain. Because residue correspondences are recomputed independently for each pair, the resulting score is not constrained by the fixed residue correspondence of the DALI MSTA, although the structures remain in the common DALI-derived coordinate frame.

**S6. Candidate retrieval, network construction, and computational benchmark**

For each target, the voxel-fingerprinting stage retrieved the top $k$candidate neighbours, which were then rescored using the pairwise geometric similarity measure described above. The $k$highest-scoring neighbours were retained, producing a directed weighted nearest-neighbour graph. We used $k=200$, based on calibration tests in which the 200 candidates retrieved for each target contained >95% of the top-30 neighbours identified by exhaustive pairwise geometric scoring. The top-30 cutoff was chosen from the observed pairwise score distributions, which levelled off around this point, providing an empirical separation between the higher-scoring neighbourhood and the lower-scoring tail. Recall was calculated as the proportion of these top-30 pairwise-scored neighbours recovered within the 200 retrieved candidates.

The graph is symmetrized by retaining the union of directed edges: an edge between targets $i$and $j$is retained if either target occurs among the other's top $k_{1}$neighbours. When an edge occurs in both directions, the higher of the two directional similarity scores is retained as its weight. The resulting sparse, undirected weighted graph forms the input to community detection. The network therefore represents pairwise structural similarity derived from independently determined residue correspondences, rather than similarity inherited from the DALI MSTA or from the voxel pre-filter.

The computational cost of this procedure was evaluated for the Worked Example 2 dataset comprising N=6,708 target proteins. With k=200 candidates per target, the pipeline performed 1,341,600 pairwise geometric evaluations. To provide a reproducible single-threaded baseline, pairwise scoring was executed serially (max_workers=1) on a virtualized cloud environment using a single thread of an AMD EPYC Milan processor. Candidate indexing required 1.00 s, while pairwise geometric scoring required 73.34 s, corresponding to an average of 54.67 μs per pairwise evaluation. The complete network-construction procedure completed in approximately 75 s. Because (k) was fixed, the number of pairwise evaluations scales approximately linearly with the number of targets, in contrast to exhaustive all-against-all comparison, which scales quadratically.

**S7. Community detection and structural populations**

The symmetrized structural similarity network is partitioned into structural communities using Infomap (Rosvall & Bergstrom, 2008). Infomap identifies communities from the flow of a random walker on the weighted network by minimizing the description length of the resulting trajectory under the map equation. Unlike fixed similarity-threshold clustering, this approach incorporates relationships among connected neighbours and does not require the number of communities to be specified in advance. Infomap was run with parameter settings “—two-level –seed 42”.

Each resulting structural community is treated as a new **population** within DaliScope. The population can subsequently be represented through MSTA views, sequence profiles, Pfam annotations, three-dimensional visualization, and the Signature Motif Pipeline. Community detection can also be applied recursively to an individual structural community to determine whether it contains finer structural subdivisions.

Existing sequence-based classifications are not used to construct the structural network or determine its communities. Pfam family and clan assignments are instead examined after community detection to assess how the resulting structural populations correspond to established sequence-based classifications.

**S8. Evaluation of structural populations**

Structural community separation is evaluated using the independently calculated pairwise geometric similarity scores. Within-community and between-community similarity distributions are compared by random sampling of structurally paired targets. Up to 2,000 within-community pairs and 8,000 between-community pairs are sampled for each community, with between-community pairs drawn from the remaining communities. Thus, up to 10,000 pairwise comparisons are evaluated per community.

This analysis tests whether targets assigned to the same structural community have higher pairwise geometric similarity than targets assigned to different communities. Pfam family and clan annotations provide an independent sequence-based reference for evaluating the organization of the resulting structural populations.

No Pfam annotation is used during voxel candidate generation, pairwise geometric rescoring, graph construction, or community detection.

**S9. Sequence profiles and structural mapping**

Sequence profiles are generated from the fixed DALI MSTA for a selected population and view. MSTA column selection determines the sequence positions represented in the profile, while population selection determines which targets contribute sequences. Sequence logos are generated using Logomaker. Stacks with IC>0 are shown.

Because profile positions retain their DALI MSTA correspondence, sequence features identified in a profile can be mapped directly to the corresponding residues in the structures.

**S10. Signature Motif Pipeline**

The Signature Motif Pipeline identifies sequence features associated with a selected structural or functional population. Candidate positions are selected from the information-content profile of the population's MSTA-derived sequence logo; the pipeline presents the five most informative positions for interactive selection. Because these positions retain their DALI MSTA correspondence, they can be mapped directly to the corresponding structural residues and inspected in three dimensions.

The selected positions define a **miniprofile** describing a sparse, Prosite-like signature motif anchored to specific MSTA positions. Rather than searching for an unrestricted sequence pattern, the miniprofile evaluates residues at the corresponding MSTA positions and is therefore restricted to structurally equivalent sites. The selected positions are converted into a position-specific scoring model and applied to target sequences using the same MSTA positional correspondence. Unless otherwise specified, a target is classified as a signature match when its score exceeds 50% of the maximum signature score.

Signature-positive targets constitute populations that can be returned to the standard DaliScope workflow for sequence, structural, annotation, and three-dimensional inspection. A signature derived from one population can therefore be tested against the broader DALI target population while retaining its structural positional context.

**S11. Rigid-body superimposition and visualization**

DaliScope provides interactive three-dimensional visualization of refined rigid-body superimpositions for a selected population. Structures are displayed in the common DALI-derived reference frame, with residue equivalences defined by the DALI MSTA available for visual inspection. The visualization allows users to assess structural agreement within the aligned region and to identify auxiliary structural elements that extend beyond the aligned region and assess whether they adopt similar spatial conformations. This provides direct structural context for interpreting population-specific similarities and differences identified by the preceding analyses.

**S12. Implementation and data workflow**

DaliScope is implemented in Python (≥3.9). Numerical and tabular analysis uses NumPy, pandas, and SciPy. Structural network community detection uses Infomap. Sequence logos are generated with Logomaker; visualization uses Matplotlib, Plotly, py3Dmol, and PyMSAViz, with interactive notebook controls implemented using ipywidgets and anywidget. The ipympl backend is supported for interactive Matplotlib visualizations.

Structural search data are obtained from the DALI server. A user submits a query structure to DALI and records the resulting JOBID. The JOBID is provided to the DaliScope data-pack interface, which generates the self-contained data pack used for downstream analysis. The resulting data pack can be loaded into Google Colab or used in a local Jupyter workflow. DALI JOBID’s persist one week; the data packs persist one hour on the server and users should save a copy locally. The DaliScope distribution includes example data packs.

The data-pack workflow separates the initial structural search from downstream population analysis. Once loaded, the target set can be iteratively filtered into populations, represented through different views, subjected to structural community detection or sequence-signature analysis, and returned to the sequence, structural, and annotation workflows.
